# Comparative analysis of chloroplast genome between wild and cultivated *Rehmannia glutinosa Libosch*

**DOI:** 10.1101/2022.12.21.521534

**Authors:** Hao Yang, Yohei Sasaki, Conglong Lian, Liping Zhang, Lili Wang, Jingfan Yang, Shujuan Xue, Suiqing Chen

## Abstract

*Rehmanniae* radix, as a tonic Chinese medicine, has a good therapeutic effect on cardiovascular and cerebrovascular diseases and central nervous system diseases, and has a good development prospect in the international market. *R*.*glutinosa* Libosch has a large number of source varieties, and the wild and cultivated varieties are different types of *R*.*glutinosa* Libosch. There has been a lot of controversy in the formulation of scientific names for wild and cultivated types of *R*.*glutinosa* Libosch. In order to learn more about the genetic information of wild *R*.*glutinosa* Libosch and explore the phylogenetic relationship between wild *R*.*glutinosa* Libosch and cultivated *R*.*glutinosa* Libosch. In the current study, we constructed and annotated a complete circular chloroplast genome of wild *R*.*glutinosa* Libosch. The chloroplast genome of wild *R*.*glutinosa Libosch*. is 153,678 bp in length, including two inverted repeat (IR) regions of 25,759 bp, separated by a large single copy (LSC) region of 84,544 bp and a small single copy (SSC) region of 17,616 bp. The GC content of whole chloroplast genome is 37.9%. The genome contains 133 different genes, including 88 protein-coding genes, 37 tRNA genes, and 8 rRNA genes. Among them, 23 genes contained introns, 50 SSR loci and 49 long repeats were found by Microsatellite scan, and codon use showed obvious A, T base preference. Nine variation regions were screened by mVISTA analysis and combined with DnaSP analysis, and psaJ-rpl33 was finally identified as a region with high variation, which could be used as a candidate DNA barcode for later interspecific identification of *R*.*glutinosa* Libosch.. Neighbor-joining method phylogenomic analysis showed that wild *R*.*glutinosa Libosch*. formed a monophyletic group, and was sister to other groups of *R*.*glutinosa Libosch*.The results of this study revealed the chloroplast genome information of wild *R*.*glutinosa Libosch* in detail, and found the genetic information difference between wild and cultivated *R*.*glutinosa Libosch*. which laid the foundation for the variety identification, genetic engineering, biosynthesis and other studies.

## Introduction

*Rehmanniae* radix is the fresh or dried tuber root of *Rehmannia glutinosa* Libosch of Scrophulariaceae. With a long history of cultivation and medicine, *Rehmanniae* radix has been widely used in the clinical treatment of human diseases. Globally, *R*.*glutinosa* Libosch is mainly produced in China, Japan and other places. Chinese scholars Lou Zhicen, Wen Xuesen, Zhou Yanqing et al. have carried out researches on the chemical composition, cultivation and genetic diversity of *R*.*glutinosa* Libosch, completed the mechanism analysis of the main chemical composition and pharmacological effect of different varieties of *R*.*glutinosa* Libosch, improved and constantly improved the quality of *R*.*glutinosa* Libosch, and optimized the planting, cultivation and production and processing methods. Meanwhile, the genetic structure and content differences among *R*.*glutinosa* Libosch varieties and regions were analyzed[1]. Japanese scholars, represented by Hoon Kitagawa et al, have carried out analysis of rhizome components of *R*.*glutinosa* Libosch., studies on components of *R*.*glutinosa* Libosch from China, Japan and South Korea and chemical composition changes before and after processing. They have reported for the first time that fresh *R*.*glutinosa* Libosch contains iridoid glycosides and the main component is catalpa, and successfully isolated four new iridoid glycosides[2]. In order to increase yield and cultivate excellent varieties, scholars from various countries have done a lot of research on the germplasm resources of *R*.*glutinosa* Libosch, genetic breeding, cultivation and planting, chemical composition and efficacy, extraction and analysis of active ingredients, plant tissue culture, virus-free rapid propagation and molecular biology.

By summarizing previous studies on *R*.*glutinosa* Libosch, it was found that as early as one thousand years ago, *R*.*glutinosa* Libosch started to “Wild varieties to cultivated varieties”. There were obvious differences between wild *R*.*glutinosa* Libosch and cultivated *R*.*glutinosa* Libosch in terms of microscopic internal genetic information, content of active components and macroscopic external morphology and physiological characteristics[3, 4]. Both wild and cultivated *R*.*glutinosa* Libosch belong to the same plant with very similar morphological characteristics. However, there were significant differences in the size of underground tubers. Therefore, different scholars have different naming and classification of *R*.*glutinosa* Libosch. Both Chinese Flora and Chinese Pharmacopoeia unified the scientific name of *R*.*glutinosa* Libosch as *R*.*glutinosa* Libosch from the perspective of large species. According to Xie Zongwan, the quality of wild *R*.*glutinosa* Libosch is different from that of cultivated *R*.*glutinosa* Libosch because of the different active ingredient content of the underground parts. In order to ensure the correct supply according to the drug name, the two scientific names are still suitable for separate use[5], several papers reported that the genetic basis of the cultivated varieties was narrow, the genetic diversity at the genome level was low, the genetic differentiation between varieties was not obvious[6], and the wild populations of *R*.*glutinosa* Libosch had higher genetic diversity than the cultivated varieties, which could be used as the screening material for the excellent germplasm resources of *R*.*glutinosa* Libosch.

With the continuous development of molecular biology technology and the continuous innovation of high-throughput sequencing technology, researchers can explore and study from the level of the whole genome of organisms[7]. Currently, the whole genome sequencing of cultivated *R*.*glutinosa* Libosch has been completed and the genetic diversity and function of some genes have been explored[8], but there are few background reports on the genetic information of wild *R*.*glutinosa* Libosch. Due to its high conservation and strong species resolution, chloroplast genome has gradually become a potential super barcode for near source inter-species and intra-species identification within the genus[9]. Therefore, this study assembled the complete chloroplast genome of wild *R*.*glutinosa* Libosch based on the next generation sequencing technology, and analyzed the basic structure of the chloroplast genome of wild *R*.*glutinosa* Libosch. The phylogenetic relationship between wild *R*.*glutinosa* Libosch and cultivated *R*.*glutinosa* Libosch was revealed by constructing phylogenetic tree, which provided molecular basis for the classification of wild *R*.*glutinosa* Libosch and cultivated *R*.*glutinosa* Libosch, and also provided important reference for genetic engineering, biosynthesis and variety improvement of *R*.*glutinosa* Libosch.

## Materials and methods

### Plant materials, DNA extraction, and DNA sequencing

Fresh leaves of wild *R*.*glutinosa* Libosch were collected from Wanxianshan (113°61′78.22″E, 35°72′27.10″N) in Xinxiang City, Henan Province,China and stored in the herbarium of Henan University of Chinese Medicine with voucher specimen number:HZYYHC15. Total genomic DNA was extracted using a Rapid Plant Genomic DNA Isolation extraction kit. The concentration of total DNA was detected quantitatively by fluorescence meter Qubit and the quality of total DNA was detected by 1% agarose gel electrophoresis.This experiment adopts the Illumina HiSeq PE150 platform, a 300 bp (insertion size) double-ended library was constructed by splicing DNA.

### Genome assembly, annotation, and analysis

Quality control software (fastqc) is used to conduct quality control on raw data and quality assessment before and after quality control. Trimmomatic software was used to cut the low-quality regions in Illumina sequencing data to obtain relatively accurate and effective data. SPAdes was used to assemble the obtained cleanreads after quality control to obtain contigs. Appropriate chloroplast contigs were selected according to the similarity and coverage between sequences, and then GapFiller software was used to fill the vacant sites of each chloroplast genome contigs. Finally, sequence corrections are used in a PriSeS-G, to correct the editing errors[10]. and insertion and deletion of small fragments in the splice process, to obtain the final complete chloroplast genome sequence. The Fasta file of the chloroplast genome obtained in the above step was uploaded to the online analysis software GeSeq (https://chlorobox.mpimp-golm.mpg.de/geseq.html#), and the IR reverse repeat region and rps12 trans-shear gene were selected to annotate. tRNA gene was annotated by tRNAscan-SEv2.70 software[11]. and wild *R*.*glutinosa* Libosch sequence published in NBCI (MW013795) was selected as the reference sequence, and the whole chloroplast genome physical map was automatically generated from the output results. Meanwhile, software MEGA 7.0 (molecular evolutionary genetics analysis 7.0) was used to analyze the chloroplast genome GC content of the sequenced samples.

### Repeat sequences, SSRs, and codon usage analysis

Random repeat analysis of chloroplast genome of wild *R*.*glutinosa* Libosch was conducted by using the functional module of REPuter of online program BiBiServ2 (https://bibiserv.cebitec.uni-bielefeld.de/reputer/) Select Forward/direct, Reverse, complementary, and Palindromic to analyze the four types of repeats. The Hamming Distance is set to 3 and the program defaults for the remaining parameters, which are 60 for Maximum Computed Repeats and 30 for Minimal Repeat Size[12]. The online program MISA (https://webblast.ipk-gatersleben.de/misa/) was used for SSR analysis of the chloroplast genome of wild *R*.*glutinosa* Liboschi, and the parameters were set as: the number of single nucleotide repeats was not less than 10, the number of double nucleotide repeats was not less than 5, and the number of trinucleotide and above repeats was not less than 4. The four, five and six nucleotide repeats were no less than 3[13]. All the protein coding sequences (CDS) of chloroplast genomes of different wild *R*.*glutinosa* Libosch were extracted, CDS with repetition and length less than 300bp were eliminated and codon bias analysis was performed on them by CodonW1.4.2 (http://downloads.fyxm.net/CodonW-76666.html). The fasta file containing CDS sequence information was input, and the out and blk files were output to obtain the number of each codon and RSCU value of all the encoded amino acids in CDS.

### Comparative analysis of chloroplast genomes

Using mVISTA (http://genome.lbl.gov/vista/index.shtml) online analytical tools to the wild *R*.*glutinosa* Libosch as reference sequence and the remaining 5 cultivated *R*.*glutinosa* Libosch visualization genome-wide comparison analysis[14]. DnaSP v6.0 was used to determine the nucleic acid variation values among six *R*.*glutinosa* Libosch chloroplast genomes. Midpoint and K(JC-Total) were used for Excel mapping, and mVISTA output was compared to mark the variation areas. Using on-line analysis software IRscope (https://irscope.shinyapps.io/irapp/) to upload GenBank annotation format files, generate visual boundary analysis diagram, The boundary information of the LSC region, SSC region and reverse repeat IR region of the annotated wild *R*.*glutinosa* Libosch chloroplast genome was compared, and the contraction and expansion of the IRa and IRb regions were analyzed.

### Phylogenetic Analysis

In order to further explore the genetic relationship between wild *R*.*glutinosa* Libosch and cultivated *R*.*glutinosa* Libosch,We selected 16 chloroplast genome sequences of *Rehmannia* and *Lysionotus pauciflorus* Maxim and *Boea hygrometrica* as outgroups, and these chloroplast genome sequences were compared by Alignment function of online software MAFFT[15]. The compared files were uploaded to MEGA6 software, and the system evolution tree was constructed using the maximum likelihood method[16].

## Results and Discussion

### Genome sequencing and assembly

In general, there are many incorrect or low-quality reads in the original data obtained by sequencing. In order to ensure the accuracy of subsequent analysis, quality control must be carried out on the original data. The total number of read sequences in this sequencing was 20.66 million, the total amount of data was 2.9Gb, and the average sequence length was 145bp. Bases with base detection quality above 30 accounted for 88.82% of the total number of bases, and the GC content of the original data was 38.78%. The above indicated that the error rate of the whole sequencing base was low, which met the experimental requirements. (S1 Table).The sequence was uploaded to NCBI, and the serial number was MW007380.

### Features of the wild *R*.*glutinosa* Libosch chloroplast genome

Like other angiosperms, the chloroplast genome of wild *R*.*glutinosa* Libosch has a typical tetrad structure, consisting of two reverse repeat regions IRs, a large single-copy region LSC, and a small single-copy region SSC. The total length of the splice wild *R*.*glutinosa* Libosch genome was 153,678bp (Fig 1). and the length of each segment was as follows: 84,544bp(LSC), 17,616bp(SSC), 25,759bp(IRs). GC content is an important characteristic of nucleic acid sequence composition and can be used as an indicator of evolution[17].The total GC content was 37.9%, the skew of AT ranged from -0.0218-0.0058, the skew of GC ranged from -0.0471-0.0405.the use frequency of A/T was higher than that of G/C, indicating that wild *R*.*glutinosa* Libosch has A/T base preference, and the highest GC content was in IR region (43.1%), followed by LSC region (35.95%) and SSC region (32.17%) (S2 Table). The results of this experiment echo those of other plants[18].

**Fig 1.**
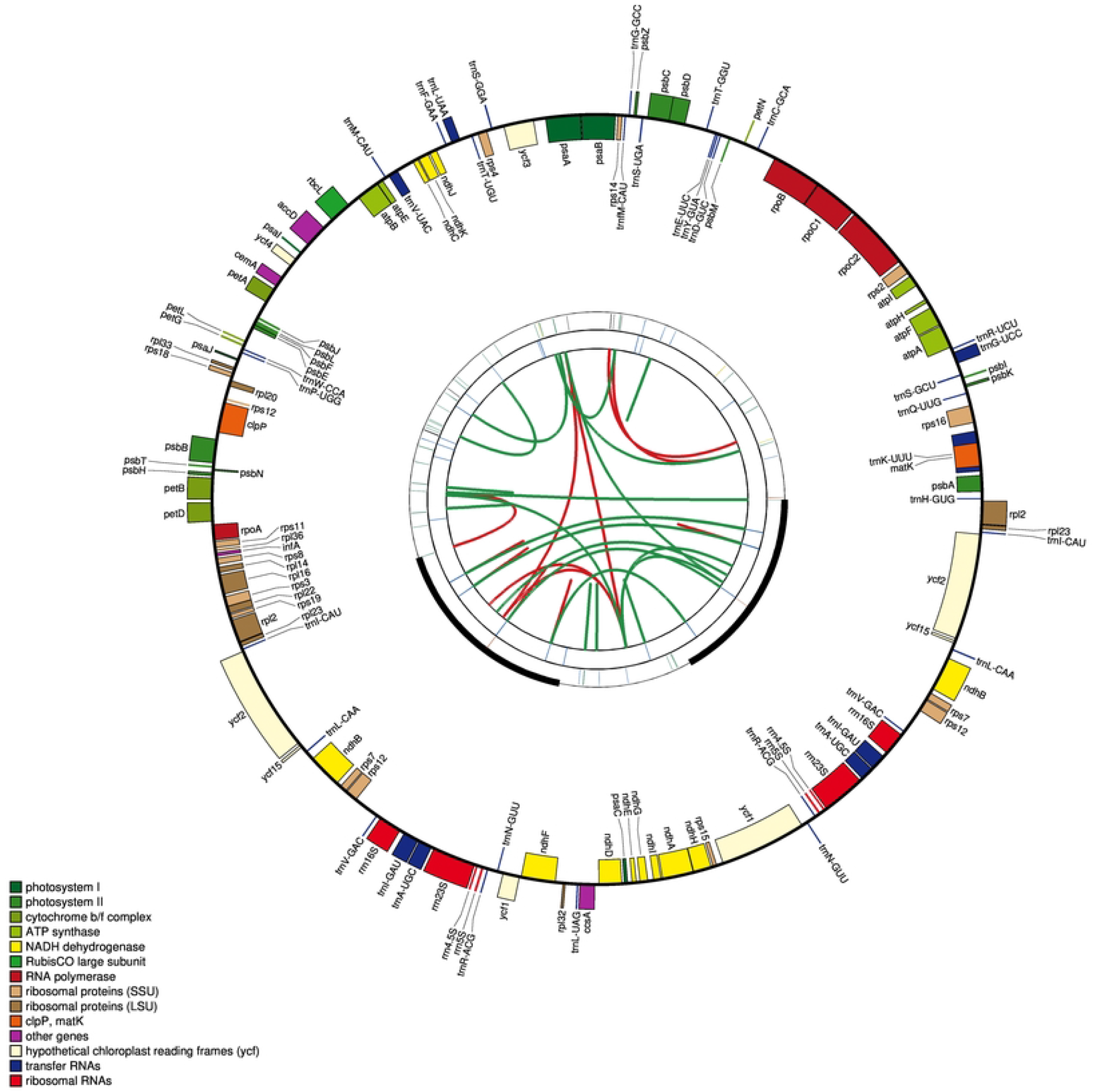
Circular gene map of the wild *R*.*glutinosa* Libosch chloroplast genome. The map contains four rings. From the center going outward, the first circle shows the forward and reverse repeats connected with red and green arcs respectively. The next circle shows the tandem repeats marked with short bars. The third circle shows the microsatellite sequences identified using MISA. The fourth circle is drawn using drawgenemap and shows the gene structure on the plastome. The genes were colored based on their functional categories.

A total of 133 genes have been annotated in the chloroplast genome of wild Rehmannia rehmanniae, including 88 protein-coding genes, 37 tRNA genes and 8 rRNA genes. For most terrestrial green plants, the functions of genes in the chloroplast genome can be divided into three main categories: the first category contains genes related to self-replication, which are annotated to the The second group is the genes related to photosynthesis. The third category is other genes involved in chloroplast biosynthesis and some genes with unknown functions, including genes related to photosynthetic system, transcription, translation, cytochrome complex, amino acid, etc[19]. Genes of the above three categories have been annotated in the chloroplast genome of wild *R*.*glutinosa* Libosch (Table 1). respectively, among which ycf1 and rps12 genes are special. There were 2 copies of ycf1 gene, but only 1 copy was properly expressed, and the other copy lost its coding function due to failure to replicate completely and became pseudogene ycf1. rps12 was the only trans spliced gene in chloroplast genome, and its 5 ‘end was located in LSC region and its 3’ end was located in IR region. This phenomenon is very similar to findings in other plants species[20].

**Table 1.**
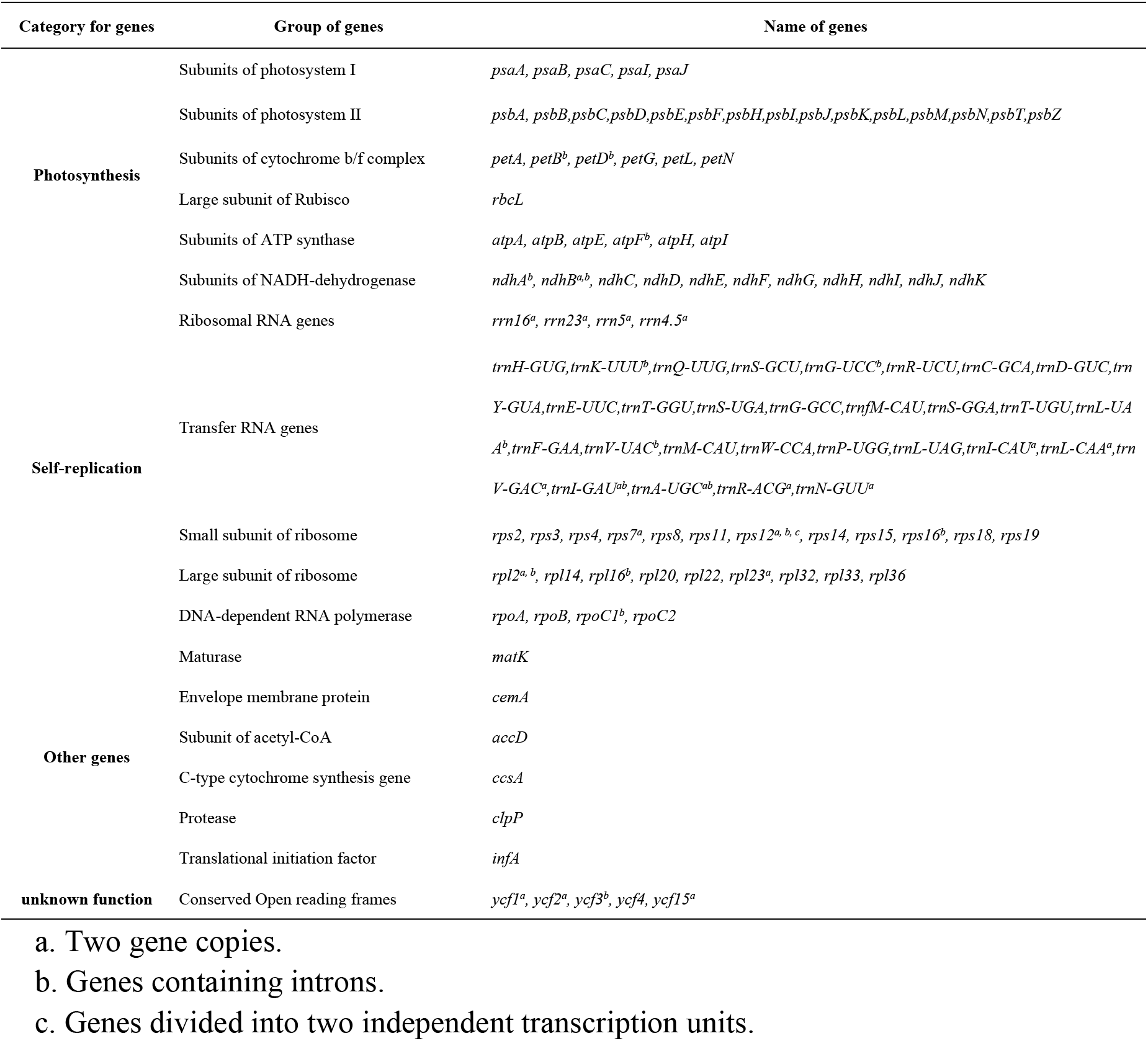
List of genes in the chloroplast genome of wild *R*.*glutinosa* Libosch.

Introns of higher plant protein-coding genes can influence gene expression activity by splicing[21]. According to the statistical analysis of chloroplast gene introns of wild *R*.*glutinosa* Libosch, 23 genes contained introns, among which rps12, ycf3 and clpP contained 2 introns, and the remaining 19 genes contained only 1 intron. (S3 Table). The statistical analysis of the length of introns and exons showed that, Most of the intron lengths in the chloroplast genome of wild *R*.*glutinosa* Libosch are between 600-900bp, and The trnK-UUU gene intron has the longest codon length of 2525, which is consistent with the results of other studies[22].At the same time, studies have shown that intron length plays an important role in gene regulation or expression[23]. Some scholars have confirmed in rice and Arabidopsis that high-expressed genes contain more and longer introns than low-expressed genes[24]. Therefore, it is reasonable to speculate that trnK-UUU may be a highly expressed gene in wild *R*.*glutinosa* Libosch.

### Codon Usage

In the long-term biological evolution, the combined effect of gene selection and mutation leads to the difference in the use frequency of most biological codons[25]. Each species has a unique codon use preference[26]. In this experiment, protein-coding genes and transfer RNA coding genes after the preliminary data screening were selected as the analysis objects, with a length of 67,445 bp. Containing 22,472 codons, the synonymous codon frequencies of different amino acids in wild *R*.*glutinosa* Libosch. chloroplast genome are significantly different (Figure 2). RSCU value is the ratio between the usage frequencies of specific codons, which is used to detect the non-uniform use of synonymous codons within the coding sequence. The codon with an RSCU value greater than 1.00 is set to 1.00, and the codon with an RSCU value less than 1.00 is lower than expected. The RSCU value of methionine (Met) and tryptophan (Trp) is 1, so there is no codon with a bias. Leucine (Leu) had the highest coding rate among the 20 amino acids, and the number of codons used was 2,353, accounting for 10.47% of the total, while tryptophan (Trp) had the lowest coding rate and the number of codons used was 419 (1.86%). The analysis of the use of four bases on the third codon of the chloroplast genome of wild *R*.*glutinosa* Libosch. showed that GC3 was 30.19%, which was similar to the GC content of the whole chloroplast genome in the previous paper[27]. further confirming that the codon of wild *R*.*glutinosa* Libosch chloroplast genome preferred to end in A and T.(Table 2).

**Table 2.**
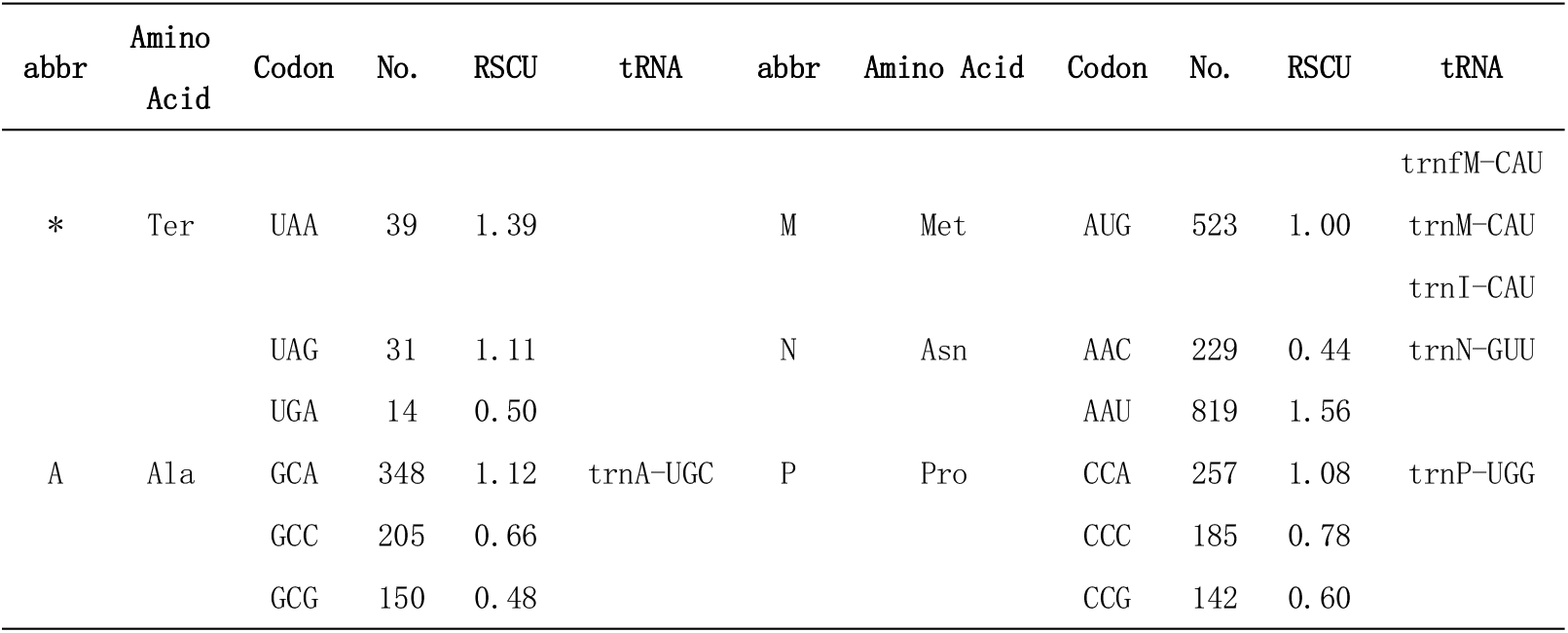
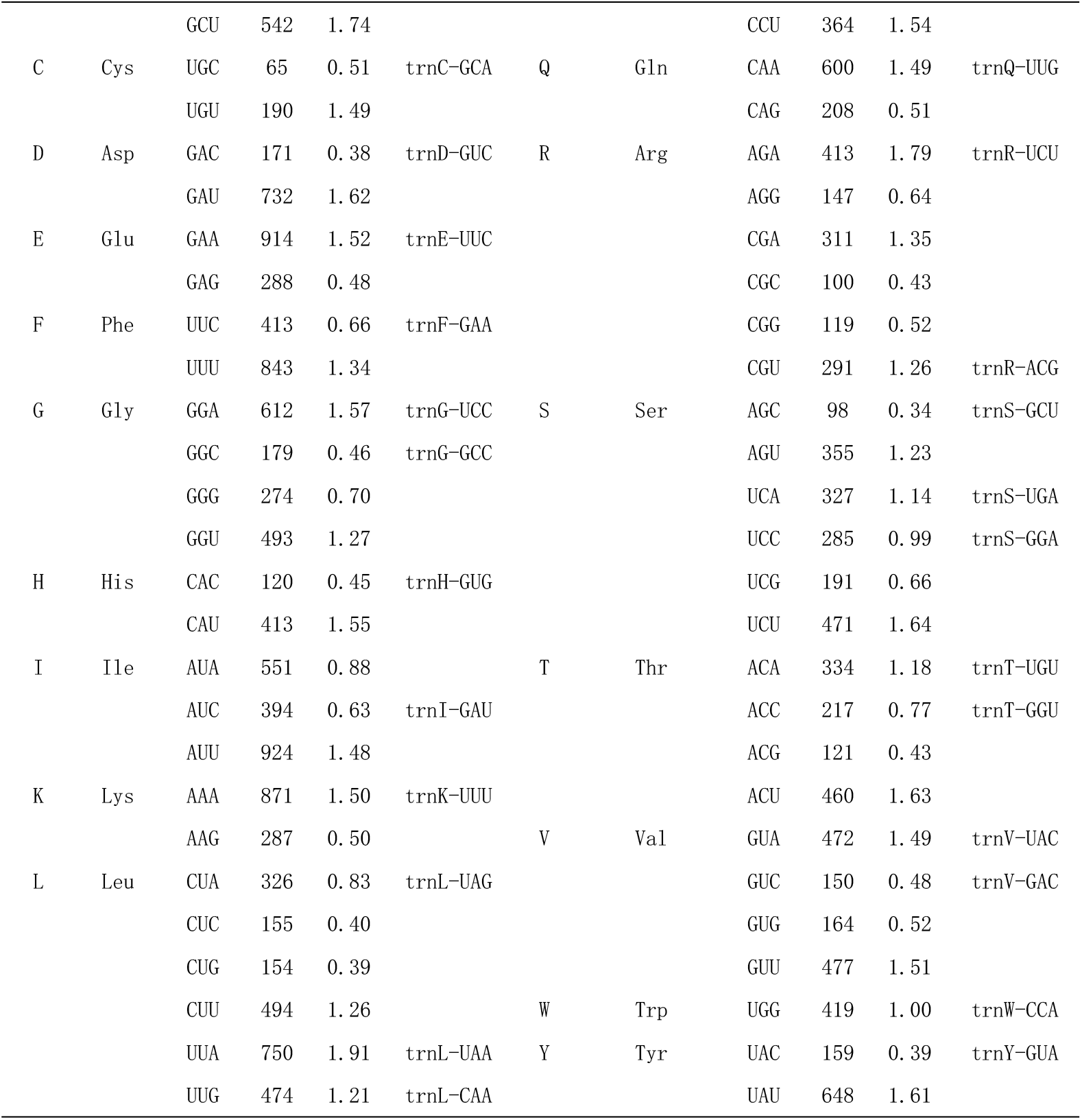
Codon-anticodon recognition patterns and codon usage in the chloroplast genome of *wild R*.*glutinosa Libosch*.

**Fig 2.**
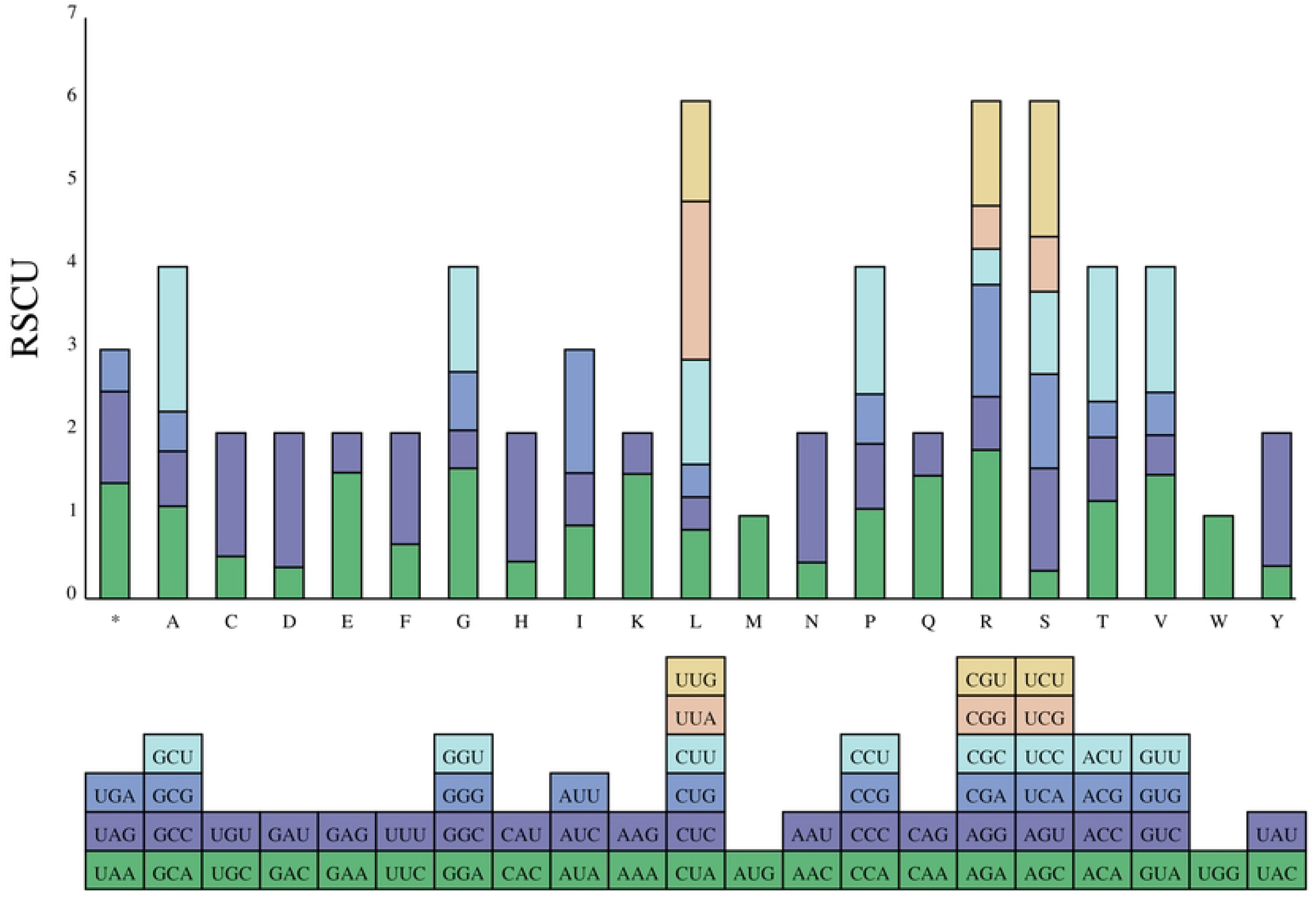
Codon bias map of the wild *R*.*glutinosa* Libosch chloroplast genome. The x-coordinate is the abbreviation for amino acid and the y-coordinate is the relative synonymous codon usage.

### SSRs and Long-Repeat Analysis

Microsatellite sequences are special sequences with highly variable length that can be distributed at different positions in the whole genome, and are widely used in species identification, population genetics and phylogenetic polymorphism studies[28, 29]. In this study, a total of 50 SSR loci were identified, including 30 single base repeats, 3 double base repeats, 2 three-base repeats, 8 four-base repeats, 3 five-base repeats and 4 complex base repeats, which were not annotated to 6 base repeats. Overall, SSRs in chloroplast genome of wild *R*.*glutinosa* Libosch have the most single nucleotide repeats and are dominated by bases A and T accounting for 60% of the total SSRs. that is, SSRs are usually composed of short A or T repeats, while few are composed of tandem G or C repeat. This is consistent with the A and T base bias in wild *R*.*glutinosa* Libosch base composition mentioned above, which may be related to the fact that A/T is more variable than G/C(Table 3).

**Table 3.**
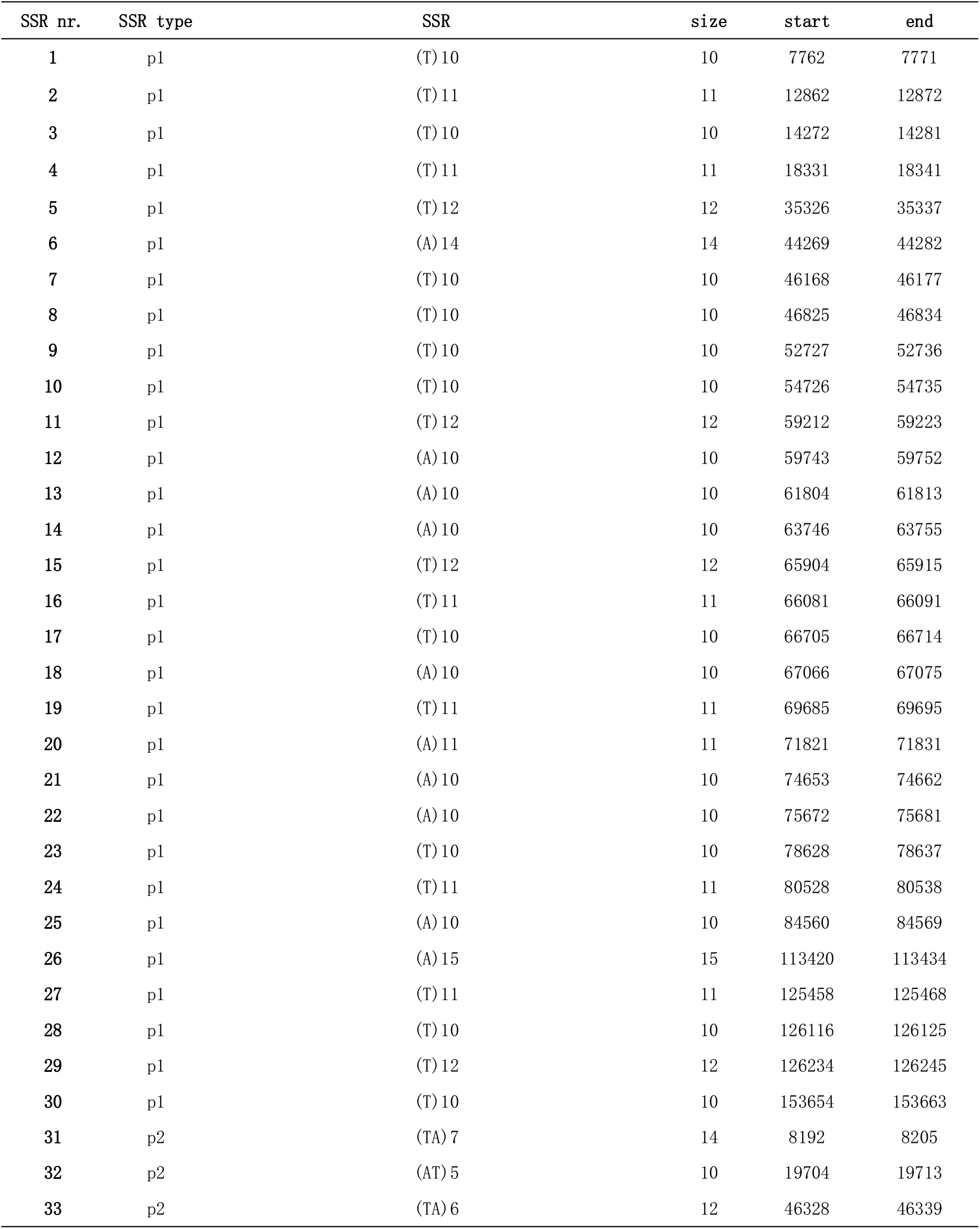
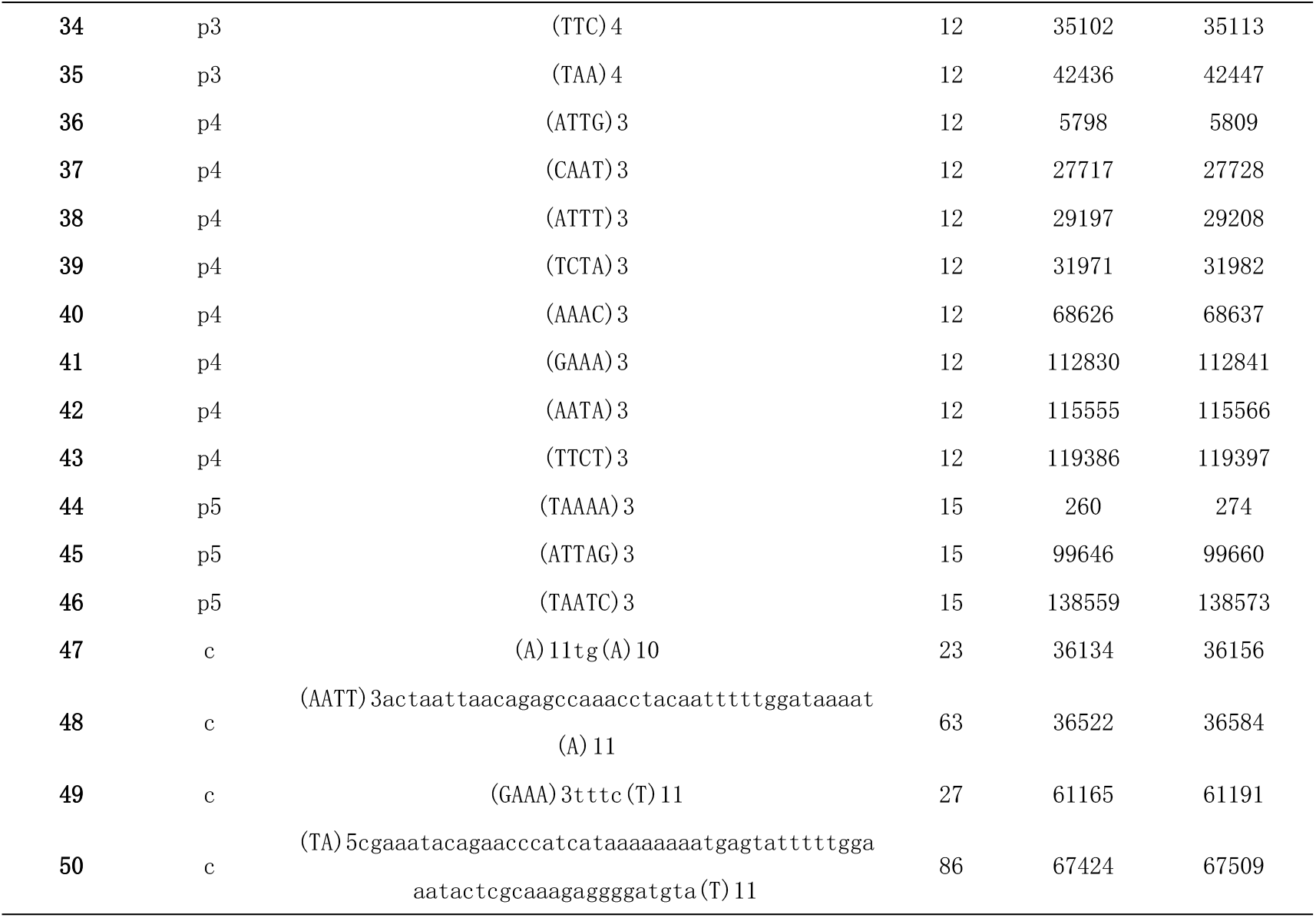
SSRs in the chloroplast genome of *wild R*.*glutinosa Libosch*.

Repeated sequences are widespread and ubiquitous in plant genomes[30]. Studies have shown that environmental changes can cause heritable changes in the number of repeats, which may imply that repeats are related to plant adaptability[31]. Long repeats include forward repeats, palindromic repeats, reverse repeats and complementary repeats. The quantitative distribution of long repeats in wild *R*.*glutinosa* Libosch showed that there were 49 repeats in total, and there were four types of repeats, mainly palindromic repeats and forward repeats, accounting for 55.1% and 34.7% respectively, and only one complementary repeat accounted for 2%. (Table 4). It was found that 3 repeats were located in the IRa region, 20 in the IRb region, 21 in the LSC region, and 5 in the SSC region. Therefore, repeats were distributed unevenly in the four regions, among which 71.4% were located in the gene coding region and 28.6% in the gene spacer region. The DNA repeats screened in this study can be used as molecular markers, genetic map construction and variety identification of wild *R*.*glutinosa* Libosch.

**Table 4.**
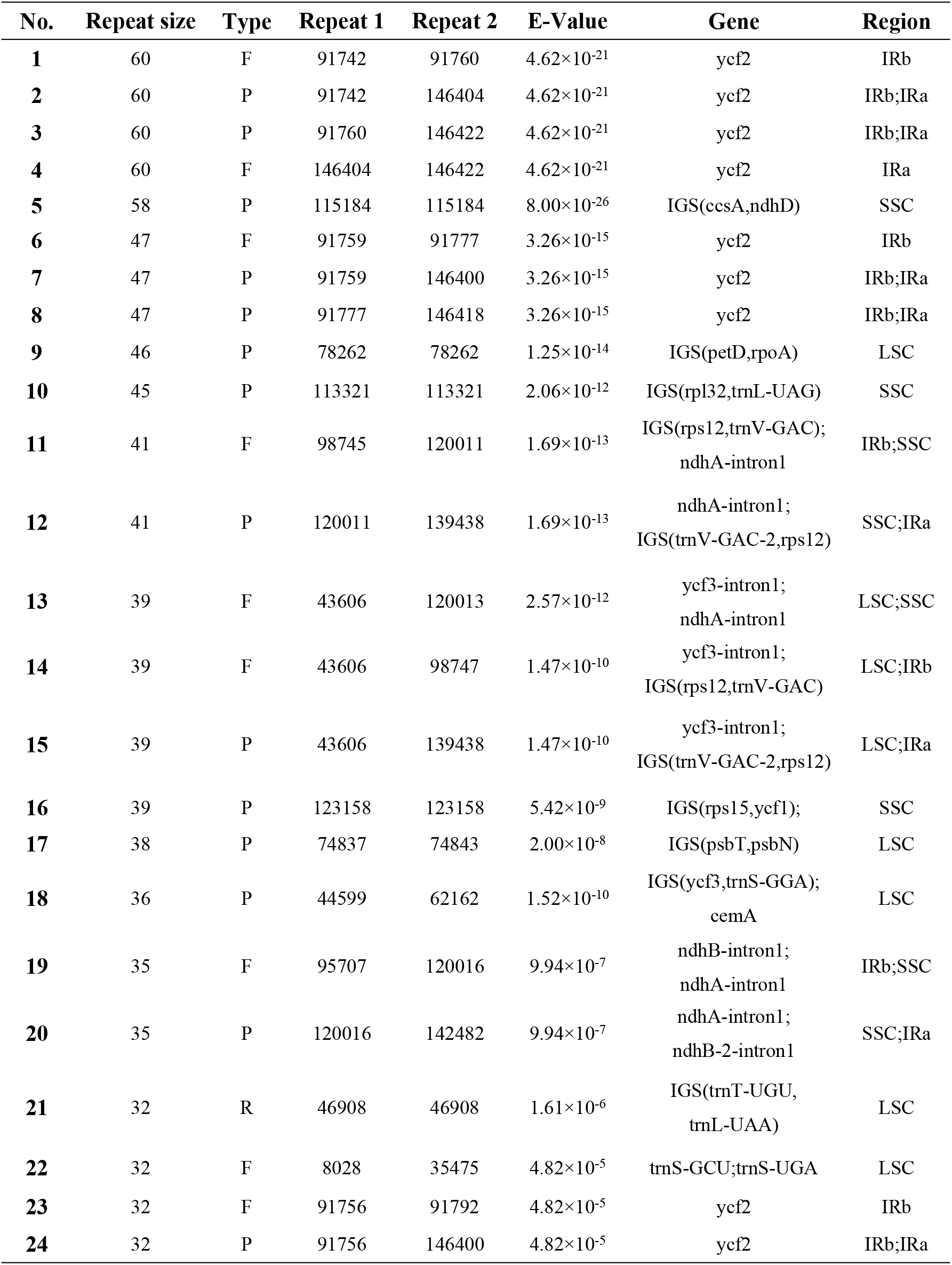
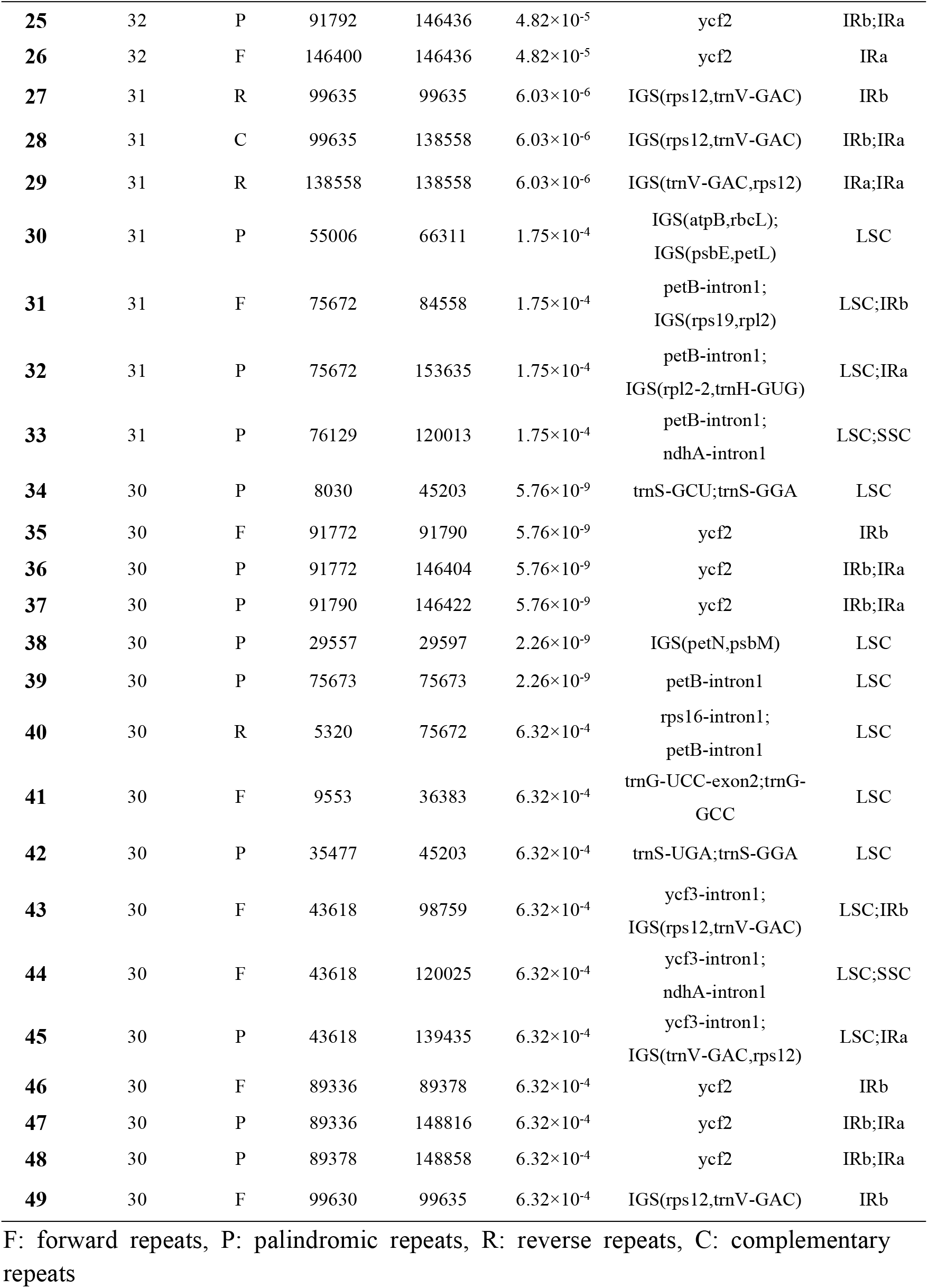
Long repeat sequences identified in the chloroplast genome of *wild R*.*glutinosa Libosch*.

### Intra-specific Comparative Analysis of Genomic Structure

Phenotypic variation of wild and cultivated *R*.*glutinosa* Libosch is obvious, and phenotypic variation can be analyzed from the two perspectives of environment and gene. There are many environmental factors and it is difficult to control them. Therefore, most scholars have carried out genetic diversity analysis of *R*.*glutinosa* Libosch with different germplasm and between different regions. However, short fragments carry less genetic information and cannot fully reflect the differences between *R*.*glutinosa* Libosch species[32]. Therefore, the chloroplast genome of wild and cultivated *R*.*glutinosa* Libosch was analyzed in this paper. The results showed that the number of genes, protein coding genes, tRNA and rRNA of wild *R*.*glutinosa* Libosch and other cultivated *R*.*glutinosa* Libosch were consistent in terms of the basic structure and composition of the genome. However, the total length of chloroplast genome of wild *R*.*glutinosa* Libosch was longer than that of cultivated *R*.*glutinosa* Libosch, and the main difference was in the LSC region. (S4 Table). According to the functional classification of the above genes, it was found that the genes related to photosystem I, photosystem II and cytochrome synthesis were mainly distributed in LSC. Therefore, it was reasonable to presume that this phenomenon might be related to chloroplast photosynthesis.

In order to further explore the size of variation between wild and cultivated species of *R*.*glutinosa* Libosch, using wild *R*.*glutinosa* Libosch as the reference genome, the chloroplast genome sequences of 6 *R*.*glutinosa* Libosch were globally compared and analyzed by mVISTA software (Fig 3). The results showed that the six chloroplast genome sequences had high similarity and basically the same sequence, among which the variation in the non-coding region (red part) was higher than that in the conserved protein coding region (blue part), the variation in the LSC region and the IR region was significantly higher than that in the SSC region and the IR region, and the rRNA gene was highly conserved without variation. It can be seen from the figure that ycf3 and ycf1 are highly variable genes, and other genes are highly conserved. Most of the genes were more than 90% similar, and the gene interregions with large variation included rpoB-trnC-GCA, psbC-trnS-UGA, psaA-ycf3, ycf3-trnS-GGA, ycf4-cemA, trnP-psaJ, psaJ-rpl33, and rps12-trnV-GAC, rps15-ycf1, These results are mostly consistent with those reported for the chloroplast genomes of other plant species[33]. these genes and regions provide strong support for further study of *R*.*glutinosa* Libosch phylogeny, genetic diversity and species identification. These regions can be used to distinguish the variation sites between wild and cultivated *R*.*glutinosa* Libosch and are candidates for barcode design.

**Fig 3.**
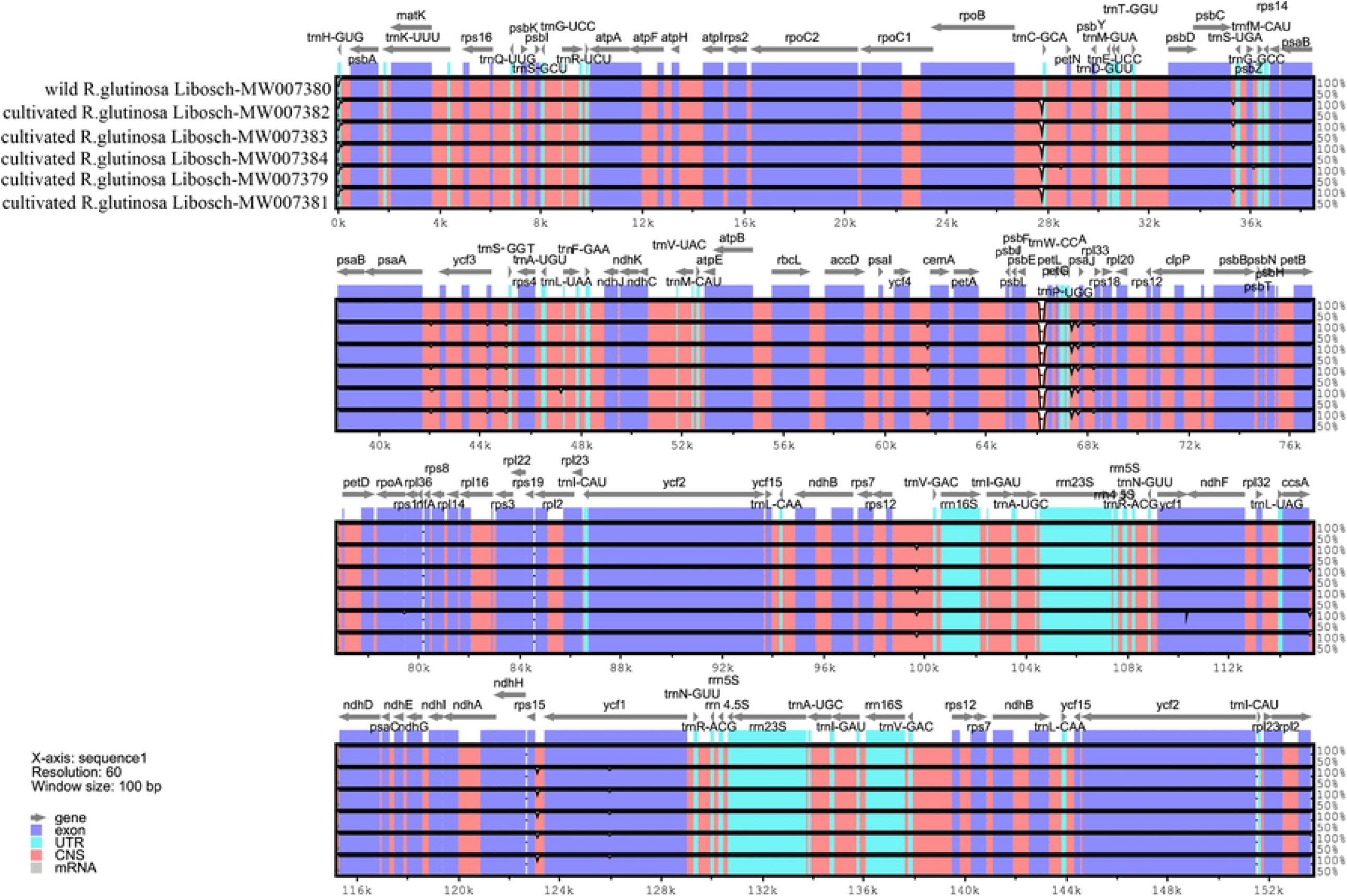
Sequence alignment of five *R*.*glutinosa* Libosch chloroplast genomes using mVISTA. the annotation of wild *R*.*glutinosa* Libosch was used as the reference. Gray arrows above the alignment indicate the position and direction of transcription of each gene. The scale on the vertical axes represents the percent sequence identity between 50% and 100%.

DnaSP software was used to analyze and detect the chloroplast genome of wild and cultivated *R*.*glutinosa* Libosch. The degree of nucleic acid variation (K value) was calculated by pairings to show the degree of variation at the sequence level. The results showed that the variation degree of the chloroplast genome IR region was significantly lower than that of the LSC region and the SSC region, and the K value was generally below 0.001. Generally, the K value greater than 0.04 was set as the highly variable gene, and the variation value within the *R*.*glutinosa* Libosch species was all lower than this value, which proved that the variation within the *R*.*glutinosa* Libosch species was small. At the same time, it can be found that there are certain nucleic acid variation between different varieties of cultivated *R*.*glutinosa* Libosch, but the variation is smaller than that between wild and cultivated *R*.*glutinosa* Libosch. Among them, five regions had nucleic acid variation values greater than 0.015, all of which were located in the LSC region, namely petN-psbM, psbD-trnT-GGU,rps14-psaB, ycf3, and psaJ-rpl33. (Fig 4). Combined with the above mVISTA analysis, Both of them were annotated to the psaJ-rpl33 region and the maximum nucleic acid variation value was 0.02384. Therefore, psaJ-rpl33 could be used as a candidate DNA barcode for later *R*.*glutinosa* Libosch variety identification.

**Fig 4.**
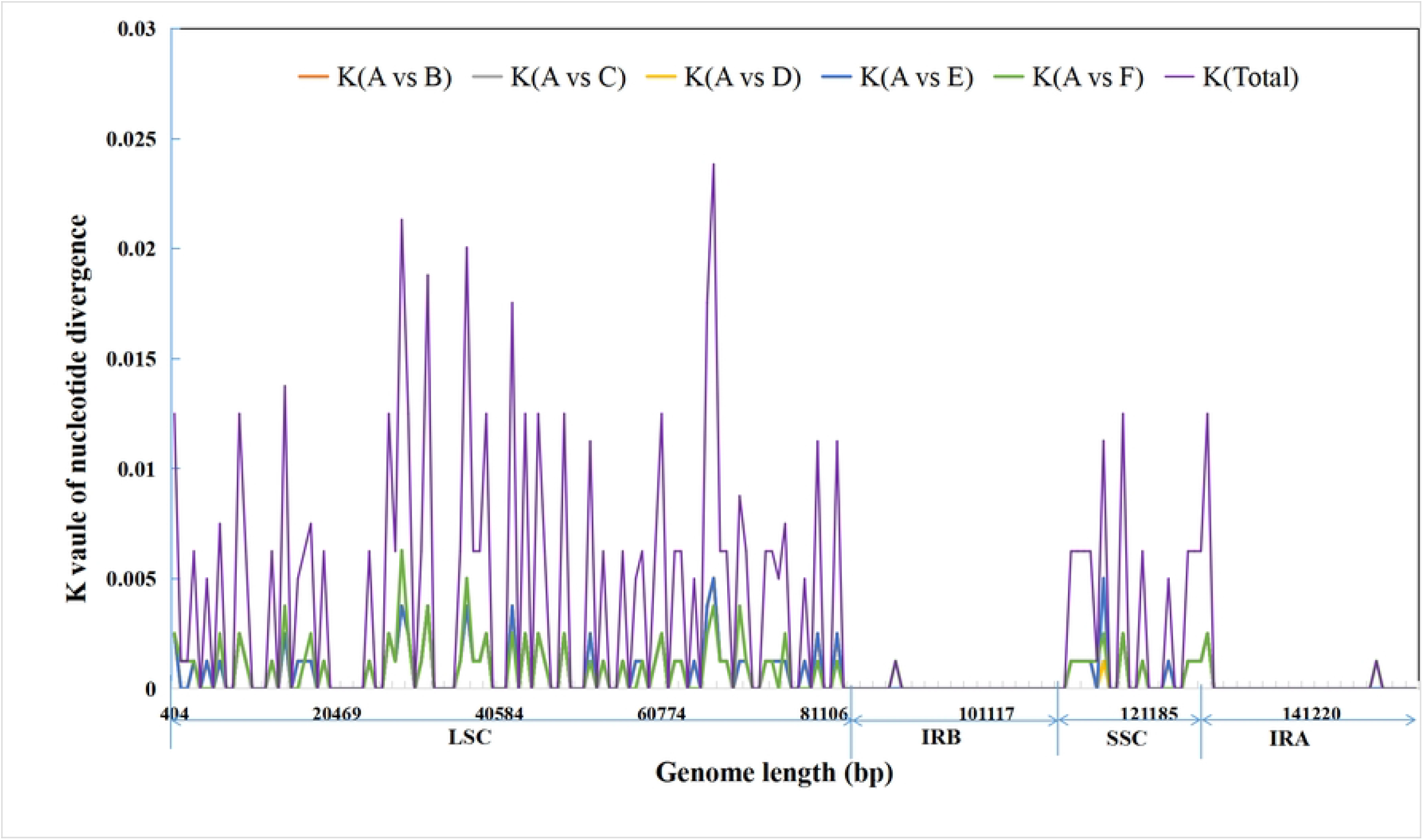
Sliding window analysis of five complete *R*.*glutinosa* Libosch chloroplast genomes. Window length: 800 bp; step size: 800 bp. X-axis: Nucleotide positions in the chloroplast genomes. Y-axis: K values of nucleotide diversity. A: wild *R*.*glutinosa* Libosch from XinXiang, Henan province (MW007380), B: cultivated *R*.*glutinosa* Libosch (MW007382), C: cultivated R.glutinosa Libosch (MW007383) D:cultivated *R*.*glutinosa* Libosch (MW007384) E: cultivated *R*.*glutinosa* Libosch (MW007379) F: cultivated *R*.*glutinosa* Libosch (MW007381) Total: the total nucleotide diversity value of A vs B, A vs C, A vs D, A vs E and A vs F.

The different size of chloroplast genome is mainly caused by the expansion and contraction of the boundaries of the four structures to different degrees[34]. We conducted a comparative analysis of the IR boundaries of the whole genome of the 6 kinds of *R*.*glutinosa* Libosch in this study. The results showed that: The JLB(LSC/IRb), JSB(IRb/SSC), JSA(SSC/IRa) and JLA(LSC/IRa) boundary genes of 6 *R*.*glutinosa* Libosch were completely consistent. In this study, the JLB boundary was rps19 and rpl2 genes, the JSB boundary was ycf1 and ndhF genes, and the JLA boundary was rpl2 and trnH genes. The JSA boundary is two genes, ycf1 and trnN (Fig 5), among which ycf1 gene extends to overlap with IRa boundary, and the overlap length range is 1083bp. These overlapping fragments made ycf1 gene have a pseudogene fragment of ycf1 at the IRa/SSC boundary. Meanwhile, it was found that wild *R*.*glutinosa* Libosch ycf1 gene had gene contraction at the SSC/IRa boundary compared with other 5 cultivated *R*.*glutinosa* Libosch species. The length of IR in the SSC region was 6bp shorter than that in other genomes. Therefore, the overall IR length of the six *R*.*glutinosa* Libosch was highly conserved, but there was still a slight difference in the boundary region between wild *R*.*glutinosa* Libosch and cultivated *R*.*glutinosa* Libosch.

**Fig 5.**
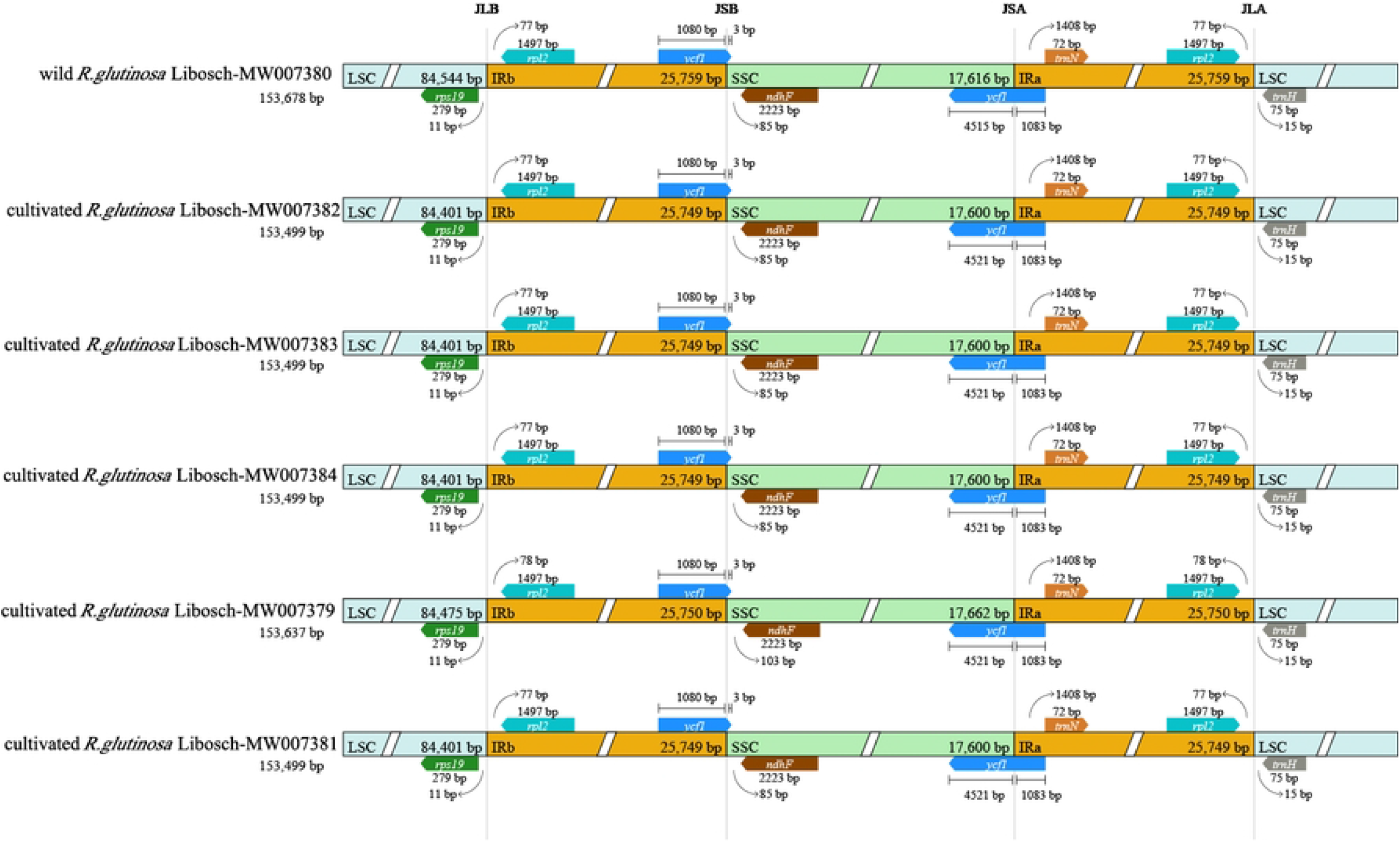
Comparison of the LSC, SSC, and IR border regions among the six *R*.*glutinosa* Libosch chloroplast genomes of examined in this study.

### Phylogenetic analysis

Chloroplast genome is of great significance for phylogenetic studies[35]. In this study, the chloroplast genome sequences of 10 *Rehmannia genus* and other groups downloaded from GenBank and the chloroplast genome sequences of 6 *R*.*glutinosa* Libosch species measured in this study were used as the features-based construction method, The maximum likelihood method (ML method) was used to construct the phylogenetic tree, in which *Lysionotus pauciflorus* Maxim and *Boea hygrometrica* were exotaxa. In the resulting phylogenetic tree bootstrap (BS) analysis showed that 11 nodes were strongly supported with BS values ≥ 90%, and 8 nodes had BS values of 100% (Fig 6). As can be seen from the results, the wild *R*.*glutinosa* Libosch measured in this study was first clustered into a single branch with 100% support rate, and then clustered into a branch with 100% support rate with 5 cultivated *R*.*glutinosa* Libosch self-sequenced and NCBI downloaded *R*.*glutinosa* Libosch, and finally clustered into a branch with Rehmannia species downloaded from GenBank database. Therefore, it can be clearly seen that there are some differences between wild *R*.*glutinosa* Libosch and cultivated *R*.*glutinosa* Libosch, which may be due to the change of gene structure in the long-term evolution process, which is consistent with the phenomenon of base loss in PsbA-trnH of wild and cultivated *R*.*glutinosa* Libosch discovered by our previous research team, and the evolutionary relationship between different species in the genus *Rehmannia* can be found. *Rehmannia henryi* and *Rehmannia solanifolia* formed the basal group of the evolutionary system as sister groups, *Rehmannia piasezkii* Maxim. differentiated successively, and *Rehmannia elata*. and *Rehmannia chingii*. sister groups formed the evolutionary group. The interspecific relationship of *Rehmannia* in this study is consistent with the results of previous studies[36].

**Fig 6.**
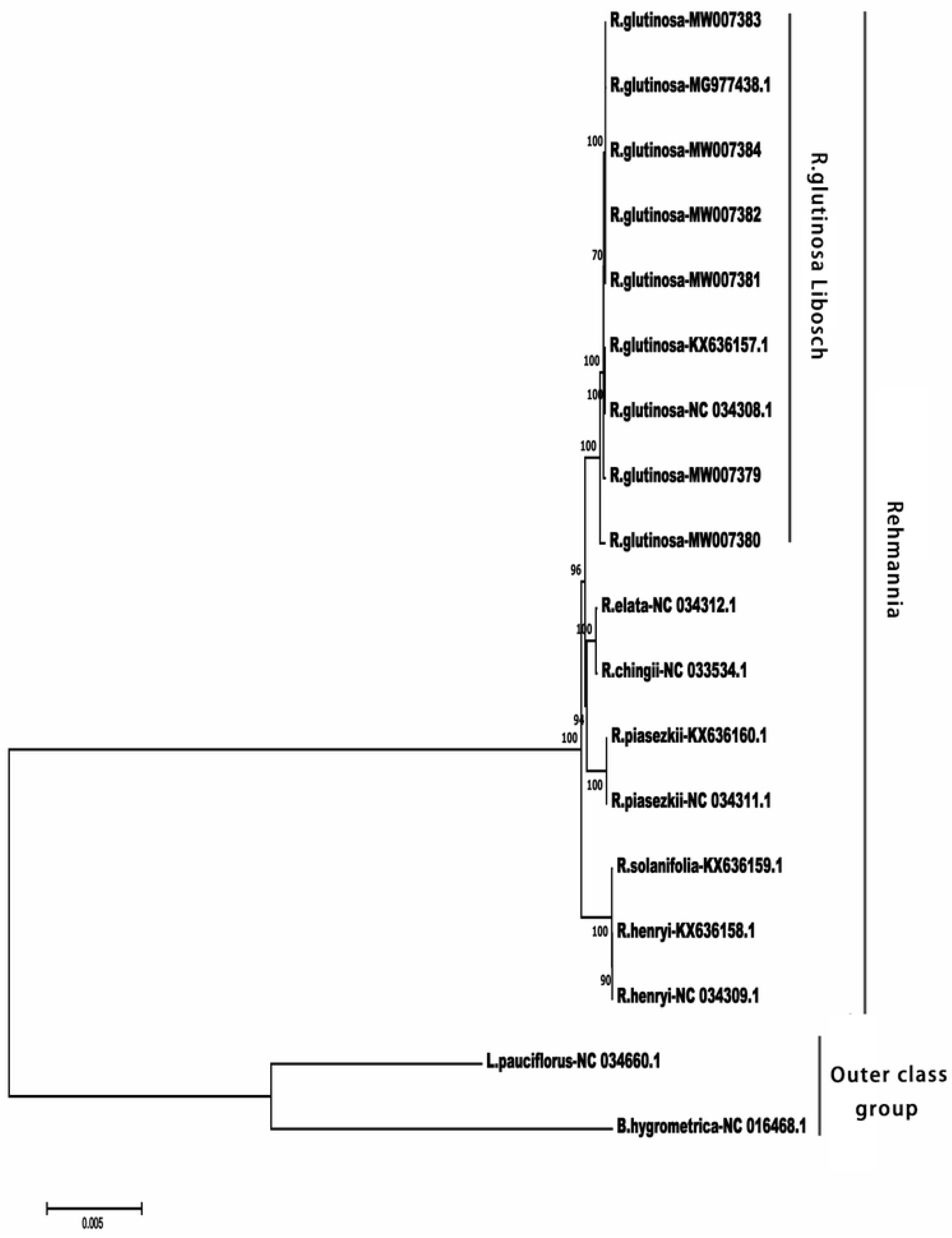
Phylogenetic tree based on the complete chloroplast genomes from Rehmannia species constructed using the maximum likelihood method. The *Lysionotus pauciflorus Maxim*.and *Boea hygrometrica*. cpDNA sequence were used as the outgroup. Bootstrap values are shown at the nodes.

## Conclusions

The comparative study of wild and cultivated products is one of the important contents of traditional Chinese medicine resources, and the accurate identification and differentiation of species is the premise and basis of biological research and resource conservation. By comparing the chloroplast genome differences between wild and cultivated *R*.*glutinosa* Libosch, this study proved that there were not only morphological differences but also significant differences in the gene level between wild and cultivated *R*.*glutinosa* Libosch. The phylogenetic tree constructed from the whole chloroplast genome can successfully distinguish wild *R*.*glutinosa* Libosch from other cultivated *R*.*glutinosa* Libosch, which provides molecular basis for the classification of *R*.*glutinosa* Libosch. At the same time, it can be seen that the chloroplast genome contains more genes and more information of loci than the previous short DNA barcode. It can make up for the lack of information loci in the combined analysis of a few chloroplast fragments, which leads to the low resolution and low support rate, and has a stronger ability of species differentiation at the level below *R*.*glutinosa* Libosch species. Therefore, the chloroplast genome has the potential as a super DNA barcode for interspecific identification of *R*.*glutinosa* Libosch.

The specificity of wild *R*.*glutinosa* Libosch should be paid more attention by relevant researchers, which is not only conducive to the research on the germplasm protection of *R*.*glutinosa* Libosch, but also has important reference value for the improvement of the quality of *R*.*glutinosa* Libosch. The continuous development of molecular biotechnology provides new technologies and new methods for this research. This study described in detail the introns, codon, repeats and other important genetic information of wild *R*.*glutinosa* Libosch. At the same time, the high variation region of psaJ-rpl33 was screened, and the DNA barcoding study of this segment could be carried out in the later stage, which not only laid a solid foundation for the smooth development of future genetic engineering and biosynthesis research of *R*.*glutinosa* Libosch. It also provided an important reference for the identification and improvement of *R*.*glutinosa* Libosch varieties.

## Acknowledgments

We are grateful to the members of the lab for their assistance and helpful discussions.

## Supporting information

**S1 Table. Raw data quality information of wild R.glutinosa Libosch chloroplast genome**

**S2 Table. Base composition of the wild R.glutinosa Libosch chloroplast genome**.

**S3 Table. The intron and exon analysis of intron contain genes in the chloroplast genome of wild R.glutinosa Libosch**

**S4 Table. Summary of six chloroplast genomes features of R.glutinosa Libosch**

